# Forest canopy, *a proxi* of light intensity, arrests *Pinus radiata* invasion: basic science to conserve the Coastal Maulino forest, Central Chile

**DOI:** 10.1101/512194

**Authors:** Persy Gómez, Maureen Murúa, José San Martín, Estefany Goncalves, Ramiro Bustamante

## Abstract

Coastal Maulino forest is an endemic forest of central Chile, which has suffered a large history of disturbance, being replaced by large extensions of *Pinus radiata* plantations. This land transformation conveys high rates of pines invasion into native remnants. In this study we examined to what extent structural features of forest patches explains invisibility of this forest-type. Within eight forest fragments, we sampled 162 plots (10 x 10 m^2^ each). We quantified seedling pine density and related this estimates with tree cover, litter depth, PAR radiation, and diversity of the resident community. Our results indicate that canopy cover was the most important variable to determine seedling pine density within forest fragments. To preserve the Coastal Maulino forest and the biodiversity containing on it, it seems to be necessary to maintain the native canopy cover. These actions can be highly effective even if we cannot avoid a massive seed arrival from pine plantations which will be unable to regenerate under well conserved native forests.

## INTRODUCTION

Annually, millions of hectares are deforested and fragmented worldwide, transforming native forests into disturbed habitats often used for agriculture and forestry practices (Schelhas and Greenberg 1996). As a consequence the land transformation often high rates of species invasion have been detected into the native remnants (Di Castri 1989; Kruger et al. 1989; Pauchard et al. 2006).

The invasibility of native forests i.e. their capacity to resist invasion, depends on different forests features. Firstly, as more diverse are native forest, they are more resistant to invasion; the reason of that is because community is saturated, resources are depleted and no more species can establish there due to interspecific competition (Pollnac et al. 2012; Relva and Nuñez 2014). Secondly, anthropogenic disturbances reduce significantly the native cover, generating luminous and dry microsites, which in turn favors the establishment of shade-intolerant exotic plant (Bustamante et al. 2003; Hobbs 2001; Laurance 1997; Parker 2001; Richardson and Brown 1986; Viana et al. 1997). The elucidation whether these forest attributes act individually or in synergy is critical to use this information for the management and control of this threat.

*Pinus radiata* is a shade – intolerant tree species original from California (USA) (Richardson and Bond 1991). Regarded as one of the most invasive species in the Southern Hemisphere (Rejmanek and Richardson 1996), this species has impacted biodiversity significantly (Higgins and Richardson 1998; Lusk et al. 2001; Rouget et al. 2001). This invasive success is concordant with disturbance regimes: (i) fire, which promotes regeneration due to serotony (Richardson and Brown 1986) and (ii) deforestation and habitat fragmentation which modifies microhabitats (Rouget et al. 4 2001). On the contrary, the native cover resists the invasion of Monterrey pine (Richardson and Bond 1991; Richardson and Higgins 1998; Richardson et al. 1994).

In this study, we examine the invasibility of the Coastal Maulino forest, an endemic forest of Central Chile (San Martín and Donoso 1996). Specifically, we assessed whether structural attributes and micro-environmental factors of forests affect *Pinus radiata* regeneration. The Coastal Maulino forest is an endemic ecosystem which has declined significantly the last decades (Donoso and Lara 1996). For instance, ca. 67% of the original forest was replaced by *Pinus radiata* D. Don between 1975 and 2000 (Echeverría et al. 2007). Currently, forest patches are embedded in a matrix of pine plantations (Bustamante and Castor 1998). Invasive process into forest fragments occurred at least from 1945 and currently, there exists approx. 10% of reproductive individuals inside forest (Gomez et al. 2011). This is an ideal scenario to expect a massive pine invasion into forests remnants.

## METHODS

Coastal Maulino forest is located in the coastal range between 35° and 37° latitude S (San Martín and Donoso 1996). The study area was located at Cauquenes, Maule Region (−72.35°; −35.97°). Topography is heterogeneous with plains, gently slopes and creeks. The climate is Mediterranean-type with mean annual temperature of 18°C and mean annual precipitation of 709 mm, concentrated mainly in the winter season (Gómez et al. 2009; Ulriksen et al. 1979). The landscape is highly anthropogenic presenting a mosaic of native forest fragments surrounded by *Pinus radiata* plantations. Dominant native tree are *Nothofagus glauca* associated with *Persea lingue* and *Gevuina avellana* in more humid habitats and with *Nothofagus obliqua* and *Nothofagus alessandrii* in drier habitats (San Martín and Donoso 1996). Evidence indicates that *P. radiata* is invading actively these forest (Bustamante et al. 2003; Bustamante et al. 2005; Gomez et al. 2012), however, we do not know which are the main drivers of this invasive process.

Our research focused on eight forest fragments dominated by *N. glauca,* which ranged from 3 to 152 ha, summing a total of 332 ha; the areal extent of the sampled zone was approx. 60 km^2^ (Figure 1). We randomly set 162 sampling plots of 0.01 ha (properly geo-referenced), inside fragments; the number of plots per fragments was set up proportionally to their area (see Supplementary material: Appendix A). We attempted that the ratio *fragment area/total area of plots,* ranged from 1 to 2, with the exception to the case of the largest one (ratio 3.46, see Fragment 8 in Supplementary Material, Appendix A. Within each plot, we quantified seedling pine abundance (response variable), species diversity and richness of native plant species, forest cover, litter width and PAR radiation. For species diversity and species richness within plots, we identified plants at species level, we recorded their abundance (in numbers), its relative abundance and finally, the Shannon-Weiner index, excluding explicitly *P. radiata.* Forest cover was photographed using a camera SONY 10.1 mega pixel, disposed on tripod, 1.20 m height over the ground. To estimate the canopy cover (in percentage), the photographs were analyzed using the SIGMA (SCAN) software. We cannot discard some contribution of adult pines (10 m height or more) to the forest cover; however, it should be negligible given the low relative abundance of this species (Media = 3%; Standard Error = 0,9%). Litter depth (cm) was assessed using a rule 30 cm-long in 5 randomly selected points within sampling plots. PAR radiation was measured using a radiometer LI-COR, Model LI-250 Light Meter USA; these measures were taken at the ground level. We tested for spatial dependence among plots, conducting a Mantel test, correlating the matrix of the geographic distance between pairs of plots with the matrix of the pine seedling density differences between pairs of plots, using the plotrix package of R.

**Figure 1.**
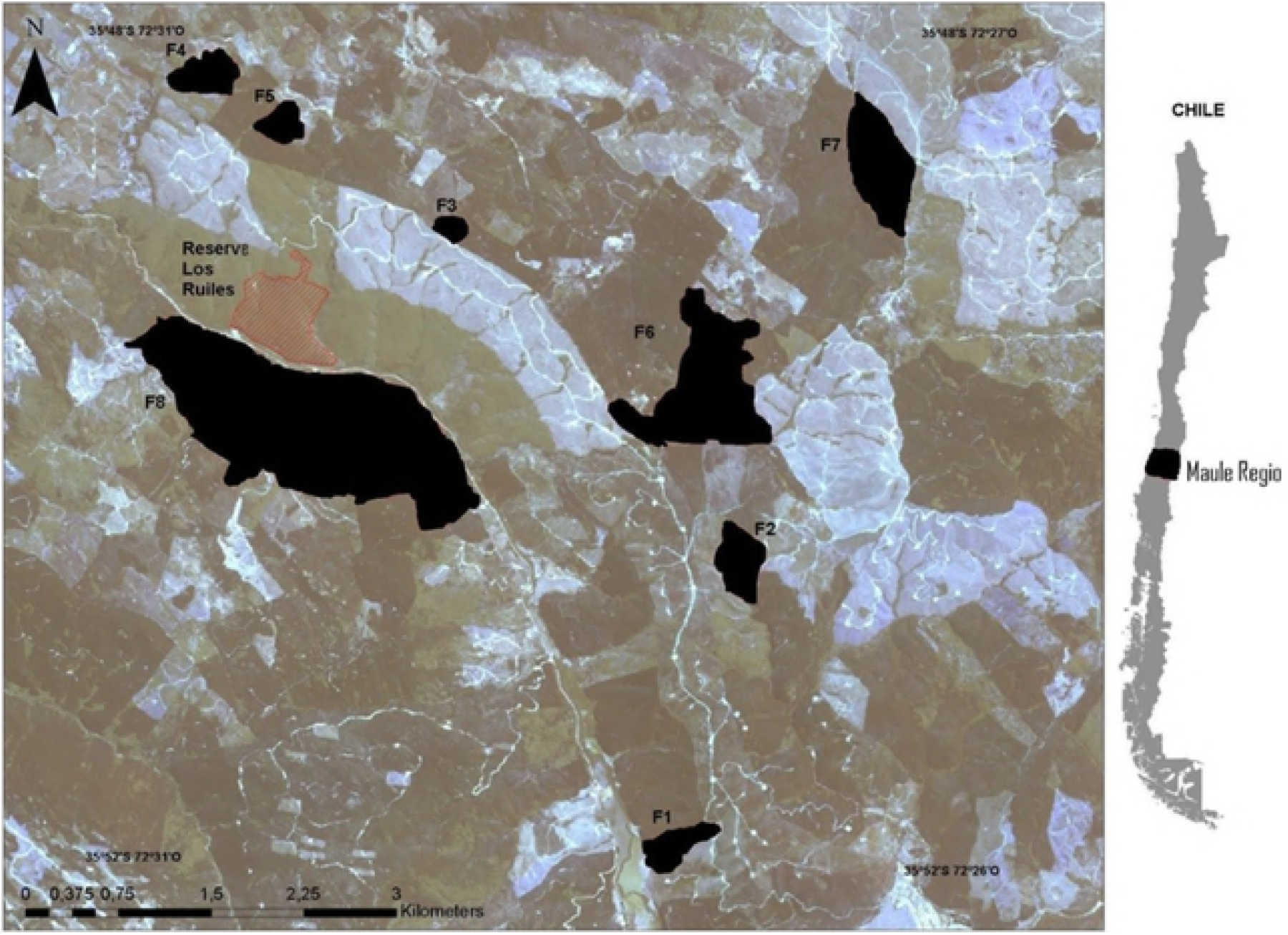
Spatial distribution of Coastal Maulino forest fragments, Cauquenes, Maule Region, Chile. Fragments of native forest remnants are shown in black (Fi = fragment i); the remaining colors indicate pine plantations in different development phases. In red: National Reserve Los Ruiles.

Within the region of study (approx. 60 km^2^, Figure 1), we found eight forest fragments, accessible for research with remarkable differences in size, isolation degree and shape (Figure 1, Appendix A). Given the extremely deforestation and fragmentation process suffered by the Coastal Maulino Forest in the past (Bustamante & Castor 1998, Etcheverría et al. 2007), it is almost impossible to selected fragments which share similar environmental conditions; then we cannot avoid to include in the analysis a suite of undesirable sources of variation, which deserve to be controlled (Figure 1). In order to do it, we considered each fragment as a block and sampling plots nested within fragments. We conducted a GLMM analysis with randomized block design (with negative binomial distribution) to assess the effect of species diversity, forest cover, litter width and PAR radiation on pine seedling density (response variable). Before the statistical analysis, we performed a Principal Component Analysis (PCA), to reduce multi-collinearity among independent variables (Graham 2003; Philipp 2001); then, the GLMM analysis was run using the fragments as randomized blocks and the two first PC as fixed effects.

We modeled the occurrence probability of pine seedling, P(O), along a forest cover gradient, We used the hierarchical logistic Huisman-Olff-Fresco (HOF) regression models (Huisman et al. 1993). This analysis is regarded an efficient method to describe species responses along ecological gradients (Oksanen and Minchin 2002; Jansen and Oksanen 2013). The HOF models were conducted using “eHOF” package in R (Team 2014). This analysis provides a set of five models for selection. The best model was selected by AIC criteria and using 1000 bootstrapping permutations. To get the goodness of fit of the model, a *Pseudo* r^2^ was also estimated (Nagelkerke 1991).

## RESULTS

Mantel test indicated that there is no spatial dependence among pairs of plots (r = 0.015; P = 0.31) i.e. the geographical distance between pairs of plots did not affect differences in seedling abundance. From PCA, the PC1 accounted 38% of the variance (eigenvalue = 1.90), and correlated negatively with tree cover (*r* = −0.53), and litter depth (r = - 0.49) and positively with PAR radiation (*r* = 0.48) (Figure 2). PC1 is an axe that represents light availability, being negative values with low light availability and positive values an increase of light availability. On the other hand, PC2 accounted 33% of the variance (eigenvalue = 1.64), and was correlated positively with Shannon-Wiener index (r = 0.65) and species richness (*r* = 0.59) (Figure 2).Thus, PC2 is an axe that represents native species diversity, being negative values with low diversity while positive values with high diversity.

**Figure 2.**
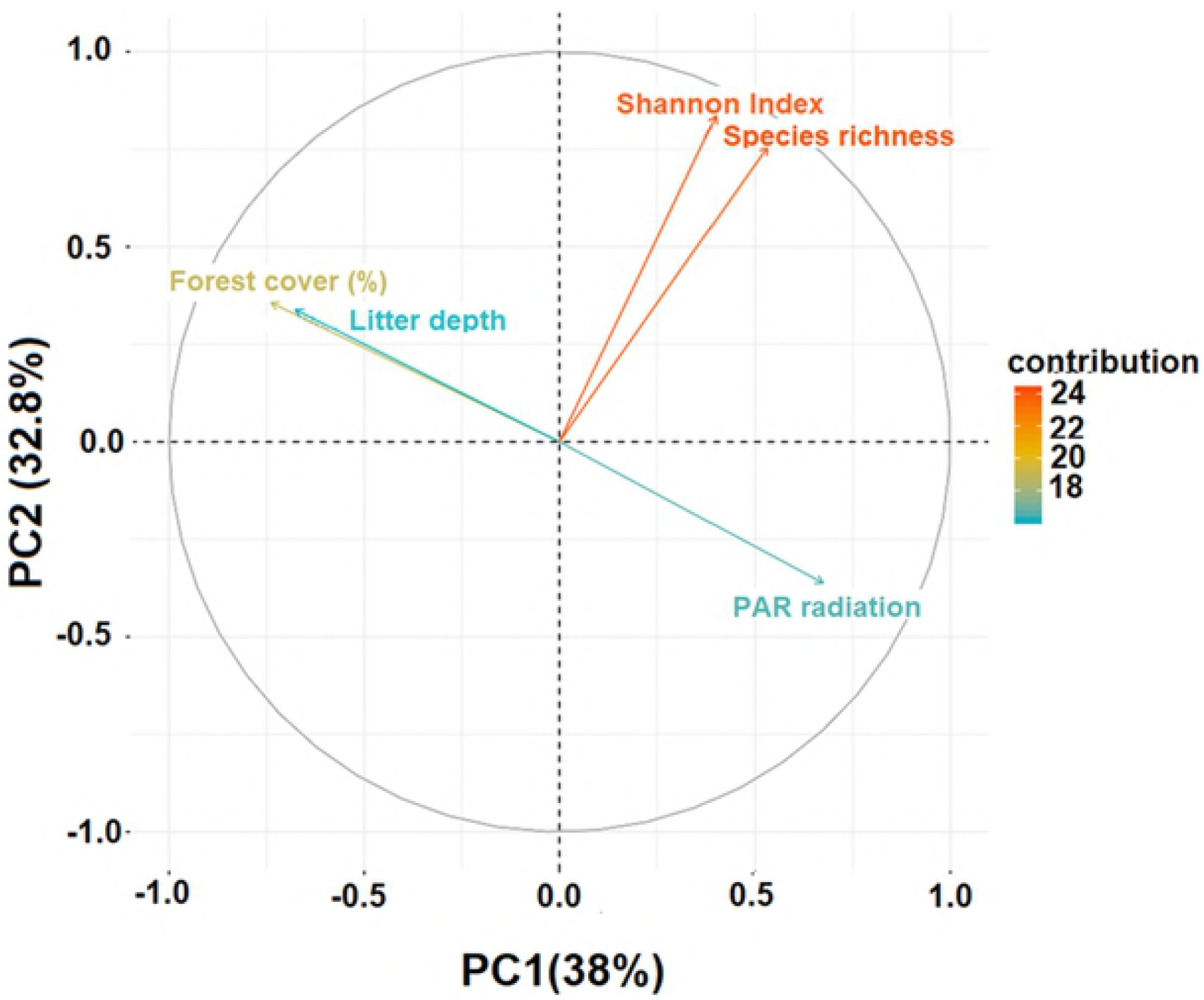
Principal Component Analysis (PCA) showing the variables that explain the data structure of independent variables related with *Pinus radiata* seedling density, Maule region, Chile. PC1: 38%; PC2: 32.8%.

From GLMM analysis, in spite of the blocks (fragments) explained an important amount of variance (0.157 ± 0.40; average ± std. dev.), we detected that pine seedling density was positively and significantly affected by PC1 (Figure 3a, Table 1); no significant effects were detected neither for PC2 (Figure 3b, Table 1) nor for PC1 x PC2 (Table 1).

**Figure 3.**
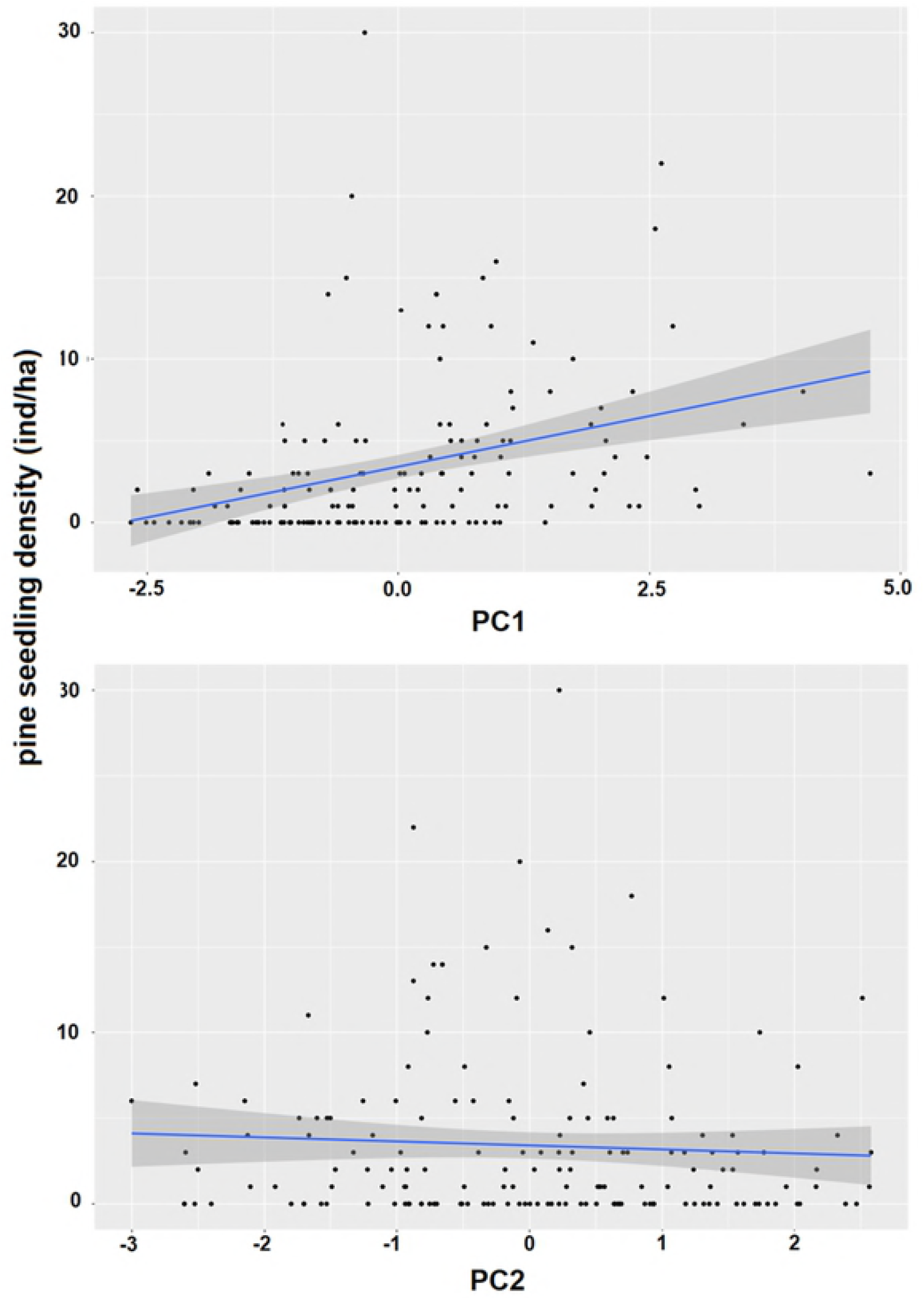
Relationship between pine seedling density and the two first Pincipal Components. a) Relationship between pine seedling density and PC1; b) Relationship between pine seedling density and PC2. For both figures, the fitted model was linear.

**Table 1.** General Linear Mixed Model (GLMM) (with negative binomial distribution), to assess the relationship between PC1 and PC2 and pine seedling abundance of *Pinus radiata* in fragmented forests, Maule Region, Central Chile.

| SOURCE OF VARIATION | D.F | Z VALUE | P |
| --- | --- | --- | --- |
| PC1 | 1 | 5,04 | << 0.001 |
| PC2 | 1 | -1,93 | 0,06 |
| PC1 x PC2 | 1 | 0,93 | 0.36 |

The HOF model-type V (Figure 4) was the best model based on its lowest deviance, AIC value and bootstrapped analysis (58%) (see Supplementary Material: Appendix B). The optimum values of P(O) occurred at a forest cover of 14%. The threshold value under which P(O) was equal to lower than 0.5 was 63%. Note that at cover values, lower than 40%, P(O) attained almost 1.

**Figure 4.**
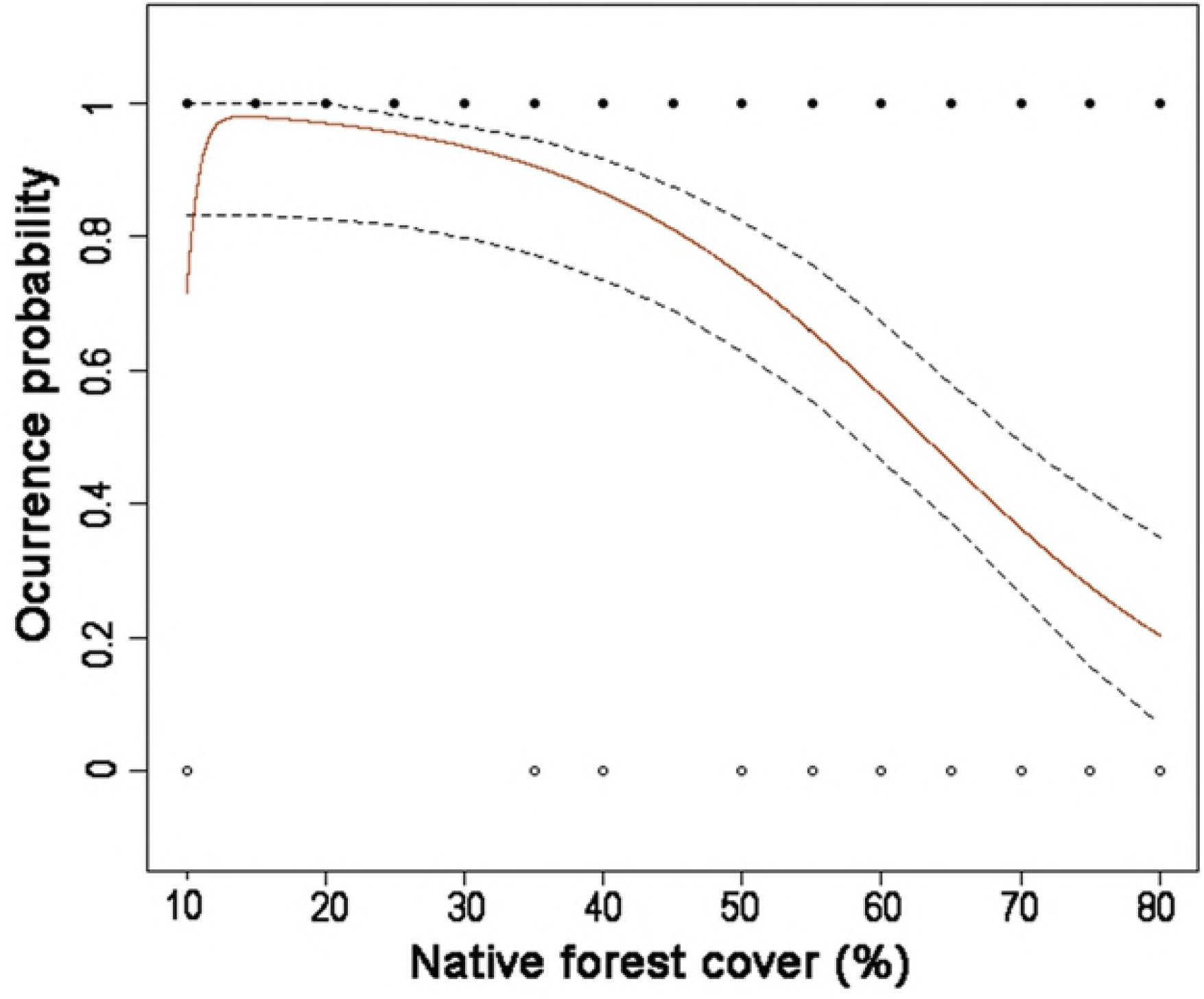
Logistic regression analysis relating forest canopy cover and Occurrence probability P(O) of *Pinus radiata* seedlings in the Coastal maulino forest. Red line describes the model; dashed lines are 95% confidence intervals. Black dots are presences; white dots are absences. For more clarity and practical use, we decided to relate P(O) with values of native forest cover directly. The analysis was conducted with n = 162 plots.

## DISCUSSION

Our results indicate that canopy cover was the most important variable to determine seedling pine abundance within forest fragments. As this variable is a proxy of light availability, as was revealed by PCA, light availability is the ultimate abiotic driver influencing pine regeneration. We did not detect direct effects of the native plant community on pine regeneration, suggesting no biotic resistance of the resident community to pine invasion. This result is of major importance because indicated no biotic filters for the colonization and establishment of this invasive tree.

Despite our study included an intense sampling effort (*n* = 162 plots) across 8 forest fragments, thus encompassing a wide spatial extent (see Figure 1), there is a notable variation of data (Figure 3) which suggests the existence of other factors, not included in our study, which may also explain pine regeneration. Several studies have tested and proven the effect of landscape variables, as for example, proximity to nearby towns and pine plantations, the forest edge extension, the size and the area/perimeter ratio on the distribution and abundance of alien plants (Bartuszevige et al. 2006; Vilà and Ibáñez 2011, Gómez et al. 2011). Future studies ought to focus a more inclusive approach encompassing landscape features in statistical models to allow a better understanding of pine invasion and improve our predictive capacity to assess invasibility of the last remnants of the Maulino forest.

The reduction of pine recruitment due to forest canopy has been also reported in other *Pinus* species such as *Pinus contorta, Pinus halepensis* and *Pinus brutia* (Richardson and Bond 1991; Richardson et al. 1994; Simberloff 2002). Based on the percentage of native cover, there have been some attempts to classify ecosystem invasibility, being forests with a very closed canopy the most resistant, followed by open woodlands, shrublands, and grasslands (Sarasola et al. 2006).

As the Coastal Maulino forest is a deciduous forest-type (San Martín and Donoso 1996), most of native trees lose their leaves during autumn-winter (like the dominant tree *Nothofagus glauca*); this fall occurs in relative synchrony with seed dispersal season of *Pinus radiata,* thus providing a temporal window with open conditions suitable for a successful seedling recruitment (Bustamante and Simonetti 2005). The high invasiveness of *P. radiata* results, in part, from its capacity to germinate (but not recruit) under a wide range of conditions (Richardson and Higgins 1998; Richardson et al. 1994, Bustamante & Simonetti 2005). If germinated seeds can survive to the next leaf fall period, when light availability increases again, then pine invasion to the interior of the forest remnants will be a matter of time (Baker and Murray 2010).

From logistic regression analysis, a reduction of the canopy cover dramatically increases the seedling density of *P. radiata,* with a threshold easy to achieve due to human disturbsnces. In consequence, any disruption of canopy cover will provide a suitable scenario for *Pinus* invasion. As human activities related to logging or recurrent fires reduce canopy cover, there are more opportunities to *P. radiata* invasion and probably other invasive species with similar regeneration requirements (*e.g. Teline monpessulana,* Gómez et al. 2012). It deserves special mention that during the last years, strong fires have occurred across Central Chile (Garfias et al. 2012). Surely, all of these fires were originated by people with devastating ecological effects (CONAF 2015; Úbeda and Sarricolea 2016). From the point of view of *Pinus radiata,* this scenario reduced strongly the native cover, therefore there are optimal conditions to increase invasion as it has been documented by previous studies in other regions of the world (Gómez and Hahn 2017; Richardson et al. 2000).

Successional studies predict successional changes of Coastal Maulino forest from a deciduous-type forest (current) to a more sclerophylous-type with a dominance of species such as *Cryptocarya alba, Lithraea caustic,* mid- and late successional tree species with perennial leaves (Bustamante et al. 2005). These successional studies did not include *P. radiata* as a novel component of the forest. Up to date, the contribution of pines to the canopy results negligible (0,3). The inclusion of pines should reinforce the predicted successional path because pines produce a dense and persistent canopy, favoring to shade-tolerants (San Martín and Donoso 1996), specially, when we know that the regeneration of *Cryptocarya alba* (a shade-tolerant tree) is not limited with *P. radiata* litter (Guerrero and Bustamante 2009).

## CONCLUSION

Our results provide basic information to conduct simple conservation actions in the Coastal Maulino forest to prevent pine invasion. To preserve this valuable ecosystem and the containing biodiversity, it is mandatory to conserve the canopy cover beyond the detected threshold of 63%. These solely actions will be highly effective even if there exists a massive seed rain that arrives each year from the surrounding pine plantations, as they are unable to recruit under well conserved native forests (Bustamante and Simonetti 2005).

## ACKNOWLEDGEMENTS

This work was supported by Anillo Project “The structure of mutualist networks in fragmented forests”, PBCT ACT34/2006, Chile. DI (Dirección de Investigación), Talca University gave partial support to the senior author. RO Bustamante was supported by ICM P05–002 and PFB-23.

## SUPLEMENTARY MATERIAL

**Appendix A.** Number of sampled plots within eight forest fragments in Coastal Maulino forest, Cauquenes, Central Chile.

| Fragment Forest | Fragment size (ha) | Plot Number<br>(100 m<sup>2</sup> each) |
| --- | --- | --- |
| 1 | 11 | 11 |
| 2 | 19 | 15 |
| 3 | 3 | 3 |
| 4 | 20 | 16 |
| 5 | 7 | 7 |
| 6 | 70 | 35 |
| 7 | 53 | 30 |
| 8 | 152 | 45 |
| TOTAL AREA | 332 | 16200 m<sup>2</sup> |

**Appendix B.** Deviance, AICC values and bootstrap tests for the selection of the best model from hierarchical logistic Huisman-Olff-Fresco (HOF) regression models (Huisman et al. 1993).

| Model | Deviance | AICC | Bootstrapped<br>(probability of the best model) |
| --- | --- | --- | --- |
| I | 215.58 | 217.61 | 0 |
| II | 179.117 | 183.19 | 0,26 |
| III | 176.286 | 182.44 | 0,06 |
| IV | 175.119 | 181.27 | 0,08 |
| **V** | **172.09** | **180.34** | **0,56** |
| VI | 175.12 | 183.37 | 0,01 |
| VII | 174.72 | 185.10 | 0,03 |

